# QTL Mapping for Pod Quality and Yield Traits in Snap Bean (*Phaseolus vulgaris* L.)

**DOI:** 10.1101/2024.04.30.591614

**Authors:** Serah Nyawira Njau, Travis A. Parker, Jorge Duitama, Paul Gepts, Edith Esther Arunga

**Author notes:** Correspondence: Edith Esther Arunga. These authors contributed equally to this work (Serah Nyawira Njau, Travis A. Parker).

## Abstract

Pod quality and yield traits in snap bean (*Phaseolus vulgaris* L.) influence consumer preferences, crop adoption by farmers, and the ability of the product to be commercially competitive locally and globally. The objective of the study was to identify the quantitative trait loci (QTL) for pod quality and yield traits in a snap × dry bean recombinant inbred line (RIL) population. A total of 184 F_6_ RILs derived from a cross between Vanilla (snap bean) and MCM5001 (dry bean) were grown in three field sites in Kenya and one greenhouse environment in Davis, CA, USA. They were genotyped at 5,951 single nucleotide polymorphisms (SNPs), and composite interval mapping was conducted to identify QTL for 16 pod quality and yield traits, including pod wall fiber, pod string, pod size, and harvest metrics. A combined total of 44 QTL were identified in field and greenhouse trials. The QTL for pod quality were identified on chromosomes Pv01, Pv02, Pv03, Pv04, Pv06, and Pv07, and for pod yield were identified on Pv08. Co-localization of QTL was observed for pod quality and yield traits. Some identified QTL overlapped with previously mapped QTL for pod quality and yield traits, with several others identified as novel. The identified QTL can be used in future marker-assisted selection in snap bean.

## INTRODUCTION

Snap bean (*Phaseolus vulgaris* L.), also known as French bean, green bean, or string bean, is a class of common bean that is grown for fresh pod consumption and for processing (Myers and Baggett, 1999). Snap beans are produced in many countries, mainly in Asia, America, Europe, and Africa (Singh et al., 2018, Parker et al., 2023). In Eastern Africa, snap bean is a major export crop with over 90% of its production destined for the international market (Mkuna, 2022). Snap bean breeding objectives vary depending on the production area but pod quality and yield are critical factors influencing the adoption of new varieties by farmers and acceptance by consumers (Singh and Singh, 2015). The optimal pod characteristics vary based on the target market segment. Typically, for the canning and packaging industry, preference lies with dark green, cylindrical, straight pods that reach uniform sieve sizes upon maturation, while the fresh market sector displays a diversity of colors and shapes (Myers and Baggett, 1999).

Most traits related to pod quality and yield are quantitative in nature and, therefore, more difficult to manipulate through targeted breeding (Singh and Singh, 2015). In order to more efficiently develop improved snap bean varieties, it is valuable to identify the causal genes or genetic markers underlying pod traits (García-Fernández et al., 2024; Saballos and Williams, 2024). Mapping of quantitative trait loci (QTL) in controlled populations can be used to identify the chromosomal regions governing traits of interest, and to develop molecular markers linked to quantitative traits. These markers are particularly valuable for breeding programs targeting traits in these categories (Singer et al., 2021).

In recent decades, advancements in genetic markers, statistical methodologies, and cost-effective genotyping techniques have revolutionized QTL mapping studies (Kaur et al., 2023). Koinange et al. (1996) pioneered the genetic mapping of pod quality traits, mapping the stringless gene (*St*) to chromosome Pv02. Their study revealed a genetic co-location between pod wall fiber and pod string, although different populations suggest a lack of linkage (Hagerty et al., 2016; García-Fernández et al., 2024). In addition, Gioia et al. (2013) characterized a *P. vulgaris* ortholog of *INDEHISCENT*, termed *PvIND* (*Phvul.002G271000*), positioned in close proximity to *St*. However, the absence of significant genetic variation at the gene locus or within its 1 kb promoter region suggested that this specific sequence might not singularly regulate pod strings. Subsequent work by Hagerty et al. (2016) identified flanking markers for *St*, spanning approximately 500 kb, from coordinates 43,984,700 to 44,472,300, within the *P. vulgaris* reference genome G19833 v2.1. Most recently, Parker et al. (2022) determined that stringless snap beans feature a tandem direct duplication of *PvIND* and a retrotransposon insertion between the repeats. Isogenic revertant lines expressing strings exhibited a reversion to a single *PvIND* copy, devoid of a retrotransposon insertion, and accompanied by a notable 50-fold reduction in *PvIND* transcript abundance. Nonetheless, the involvement of additional genes in pod suture string formation remains ambiguous.

The size of the pods holds significance not only for yield but also for commercial viability, impacting consumer satisfaction and influencing pricing in wholesale and retail markets (Li et al., 2023). Like other traits associated with yield (pod weight per plant, pods per plant), pod size is a multifaceted quantitative characteristic, highly influenced by environmental factors (Campa et al., 2018). Pod morphology encompasses factors like straightness, thickness, length, cross-section shape, and color, and dictates the subsequent utilization of the product in either the fresh market or processing. Previous research has demonstrated that pod size characteristics such as pod length, pod thickness, and pod width exhibit quantitative inheritance (Yuste-Lisbona et al., 2014; González et al., 2016; Hagerty et al., 2016; Murube et al., 2020; Wu et al., 2020; García-Fernández et al., 2021; Li et al., 2023). Hagerty et al. (2016) mapped overlapping pod wall fiber, width, and thickness QTL on Pv04, and pod length on Pv09, while Murube et al. (2020) mapped pod length, pod width, and pod thickness to Pv01, Pv02, and Pv07.

Breeding for pod quality and yield traits in snap bean is a time-and resource intensive endeavour that involves generations of inbreeding and specialized equipment because of the complexity in the genetics of pod and yield traits. Therefore, identification of genomic regions and molecular markers associated with pod quality and yield traits may permit early-generation marker-assisted selection, which would not only reduce costs, but also increase precision in selection (Kelly and Bornowski, 2018). Furthermore, breeding programs benefit from information about genetic linkage between genes controlling the same or different traits, which may affect the chosen breeding method.

The objective of this study was to identify QTLs and genetic markers associated with pod quality and yield traits in snap beans. This information is key for the development of robust molecular markers which serve as plant breeding tools to increase efficiency in snap bean improvement programs.

## MATERIALS AND METHODS

### Plant materials

A biparental mapping population of 184 F_6_ recombinant inbred lines (RILs) derived from a cross between Vanilla (female parent) and MCM5001 was used for linkage mapping and QTL analysis. Vanilla is a commercial snap bean variety with white seeds, fine market class pods (6-9 mm in diameter), and high resistance to bean common mosaic virus (BCMV), halo blight (*Pseudomonas syringae* pv. *phaseolicola*), and rust (*Uromyces appendiculatus*). MCM5001 is a dry bush bean bred for resistance to BCMV and bean common mosaic necrosis virus (BCMNV) by the International Center for Tropical Agriculture (CIAT, Cali, Colombia). Its seed are brown and cream speckled. The RILs were developed through single seed descent (SSD) in an insect-free greenhouse at the University of Embu (37° 27’ E, 0° 30’ S).

### Plant growth conditions

The 184 RILs and their parents were evaluated in three field sites in Kenya: (i) Kutus farm in Kirinyaga County (37° 19’ E, 0° 33’ S; 1,279 masl), (ii) Don Bosco farm in Embu County (37° 29’ E, 0° 34’ S; 1,259 masl), and (iii) Mariira farm in Murang’a County (36° 56’ E, 0° 47’ S; 1,255 masl), as well as in a greenhouse at the University of California, Davis (121° 45’ W, 38° 32’ N). The soils for the Kutus and Mariira farms were classified as Humic Nitisols while at Don Bosco there were Nito-Rhodic Ferralsols. The three trials were conducted during the short rain season of 2022 and supplemented with irrigation. The field trials were conducted in randomized complete block designs (RCBD) with three replications, while the greenhouse experiment was a completely randomized design (CRD) with a single replicate. In the field, each RIL and the parents were planted in a single row plot, measuring 2 m long at a spacing of 20 cm between the plants and 50 cm between the rows. The fields were plowed and harrowed to achieve a moderate tilth seedbed. Di-ammonium phosphate (18-46-0) fertilizer was applied at a rate of 200 kg ha^-1^ and thoroughly mixed with soil. During flowering, the plants were top-dressed with calcium ammonium nitrate at a rate of 100 kg ha^-1^. All cultural practices were conducted to ensure that the fields were free of pests, diseases, and weeds.

### Phenotypic data collection

Eight snap bean traits were evaluated (pod wall fiber, pod string, pod diameter, pod length, pod weight per plant, pod number per plant, pod shape and pod shattering), although pod string and pod wall fiber were evaluated using more than one criterion for both the field and greenhouse conditions as shown in Table 1. Data on pod wall fiber for dry pods were collected at the R9 stage (pod maturation; Fernández et al., 1985) while fresh pods were examined at the R8 stage (pod fill). The dry pod fiber was first evaluated for presence or absence of constrictions and secondly based on a scale of 0 (no wall fiber) to 10 (full wall fiber). The fresh pods were snapped in the middle to determine the presence or absence of fibers on a scale of 0-2 (0-no fiber, 1-few fibers and 2-many fibers).

**TABLE 1.** Description of the traits evaluated in the field and greenhouse trials of snap bean parents and RILs.

| Serial No. | Trait | Growing condition | Evaluation description | Reference |
| --- | --- | --- | --- | --- |
| 1. | Pod suture string scale (PSSS) | Greenhouse | Dry pods at R9 stage, 0-10 scale 0-no string, 10-full string | Parker et al., 2022 |
| 2. | Pod suture string (PSS) | Greenhouse | Fresh pods at R8 stage, breaking the pods at the tip and pulling the string |  |
| 3. | Pod fiber scale (PFS) | Greenhouse | Dry pods at R9 stage, 0-10 scale 0-no fiber, 10-full fiber | Parker et al., 2022 |
| 4. | Pod wall fiber (PWF) | Greenhouse | Fresh pods at R8 stage, breaking pods at midpoint and observing presence or absence of pod wall fibers |  |
| 5. | Pod length (PL) | Greenhouse | Using a ruler, length measured from the tip of the pod beak to the calyx attachment site | Hagerty et al., 2016 |
| 6. | Pod shape (PS) | Greenhouse | Visual observation 0-Round, 1-flat |  |
| 7. | Pod shattering (PSH) | Greenhouse | Visual scoring 0-no shattering, 1-pods opened and twisted | Vittori et al., 2021 |
| 8. | Pod string fresh pods (PSFP) | Field | Boiling the pods at 100 °C for 30 min and pulling the string from the calyx to the tip. String length measured as a ratio of the pod string length to total pod length | Hagerty et al., 2016 |
| 9. | Pod string dry pods (PSDP) | Field | Dry pods at R9 stage, 0-10 scale 0-no string, 10-full string | Parker et al., 2022 |
| 10. | Pod fiber fresh pods (PFFP) | Field | Fresh pods at R8 stage, 0-2 scale 0-no visible fiber strands, 1-some visible fiber strands, 2-many visible fiber strands | Hagerty et al., 2016 |
| 11. | Pod fiber dry pods (PFDP) | Field | Dry pods at R9 stage, 0-10 scale 0-no fiber, 10-full fiber | Parker et al., 2022 |
| 12. | Pod shape (PSh) | Field | Visual observation 0-Round, 1-flat |  |
| 13. | Pod weight per plant (PWPP) | Field | Dividing the total weight of the pods by the number of plants |  |
| 14. | Pods per plant (PPP) | Field | Dividing the total number of pods by the number of plants |  |
| 15. | Pod length (PLF) | Field | Using a ruler, length measured from the tip of the pod beak to the calyx attachment site | Hagerty et al., 2016 |
| 16. | Pod diameter (PD) | Field | Measured by passing the pods through holes of a bean ruler manufactured by Royal Sluis®. The holes vary in sizes as follows: $>5 \leq 6$ mm (extra fine), $>6 \leq 9$ mm (fine) and $>9$ (bobby) | |

The pod string (PS) was also evaluated for both fresh and dry pods. The fresh pods were boiled in a water bath for 30 minutes at 100 °C. The pod strings were gently pulled from the calyx along the adaxial suture of the length of the detached string, measured and recorded. The fresh pod string length was then calculated as a ratio of pod suture string length to total pod length (Hagerty et al., 2016). Pod diameter (PD) was measured by passing the pods through holes of a bean pod ruler manufactured by Royal Sluis®. The holes vary in diameter sizes ranging from 5 mm to 9 mm. The pod length (PL) was measured from the end of the petiole to the tip of the pod while the pod weight per plant (PWPP) was computed by dividing the total weight of the pods by the number of plants. Pods per plant (PP) was computed by dividing the total number of pods by the number of plants.

### Phenotypic data analysis

Statistical analyses on quantitative phenotypic data were conducted in SAS 9.3 (SAS Institute 2011). The assumption of normally distributed residuals required for analysis of variance (ANOVA) was checked for the traits measured; the results indicated that all traits were normally distributed. A combined ANOVA for the three sites was, therefore, conducted using PROC GLM for the traits based on the following statistical model:

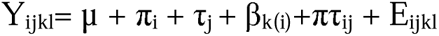

where: **Y_ijkl_**= Response variable; **µ =** Mean of the population; π**_i_** = Fixed effect of the i^th^ location; τ**_j_** = Fixed effect of the j^th^ genotype (RILs); β**_k(i)_** = Random effect of the k^th^ replication within i^th^ location; πτ**_ij_**= Fixed effect due to the interaction between i^th^ location and j^th^ genotype; **E_ijkl_** = Residual effect.

The Pearson correlation coefficient among the traits was performed using PROC CORR in SAS 9.3 version.

### Genotyping and QTL analysis

Seeds of the parents and RILs were sown in pots filled with 5 kg topsoil and placed in a greenhouse at University of California, Davis. Single nucleotide polymorphism (SNP) genotyping was accomplished by genotyping-by-sequencing (Elshire et al., 2011, Ariani et al., 2016). DNA was extracted from greenhouse-grown seeds using Qiagen DNeasy extraction kit (Qiagen, Hilden, Germany). DNA quality was checked by NanoDrop spectrophotometer and agarose gel electrophoresis. Library prep was conducted with *Cvi*AII and 150 bp paired-end sequencing was conducted on the prepared libraries at the University of California, Davis Genome Center. The reads were aligned with the v2.1 reference genome assembly of G19833 (Goodstein et al., 2013; Schmutz et al., 2014; https://phytozome-next.jgi.doe.gov/info/Pvulgaris_v2_1).

Read demultiplexing, alignment to the common bean reference genome (G19833 v2.1, (Schmutz et al., 2014), and variant calling was conducted in NGSEP3 and NGSEP4 (Tello et al., 2019, Tello et al., 2024). Data curation was performed by removing SNPs with more than 20% missing or heterozygous calls. SNPs were only kept if they had a genotype quality (GQ) over 20, were biallelic, had a minor allele frequency > 0.25, were at least 5 bp from any other SNP, were genotyped in at least 160 of 184 population members, and were found in non-repetitive regions, as defined by Lobaton et al. (2018). Individuals were plotted by missing calls and by heterozygous calls, and outliers were eliminated from further analysis. Non-parental alleles and duplicated lines were also removed. Only SNPs that were polymorphic in both the parents and the RIL population and had minor allele frequencies (MAFs) >0.25, were used for linkage mapping. After the quality checks, 5,951 SNPs were retained for linkage map construction. Linkage mapping was conducted in Rstudio using the ASMap R package (Taylor and Butler, 2017, R Core Team, 2022). QTL mapping was conducted using maximum likelihood through the EM algorithm of the R/qtl package in R (Lander and Botstein, 1989; Broman et al., 2003). A significant LOD score threshold for QTL (LOD=3.413) was developed based on the 95^th^ percentile of LOD scores of 1000 random permutations of the genotypic data. A total of 11 linkage groups corresponding to the 11 chromosomes were developed. The coefficient of determination (R^2^) was used to estimate the proportion of variation explained by a QTL (1-10^(-(2/*n*)*LOD) where *n* is the number of individuals genotyped at the locus. Results were compared with gene models located between flanking SNPs in v2.1 of the common bean reference genome (Schmutz et al., 2014).

### Candidate gene identification

The common bean genome v2.1 was browsed using Phytozome v13 to identify candidate genes in the QTL identified in this study, and data for these were extracted in Phytomine (https://phytozome-next.jgi.doe.gov/phytomine/begin.do). A gene was considered as a candidate if it was located within the confidence interval of the QTL, and its role or putative role in processes related to pod quality and yield has been established or proposed in other species.

## RESULTS

### Phenotypic analysis

#### Pod yield

The two parents (Vanilla and MCM 5001) were not significantly different in all the three sites for pod weight per plant (PWPP). The average PWPP for MCM 5001 (85.64 g) across the three sites was higher than Vanilla (75.86 g). The Don Bosco site had the highest mean PWPP across the sites while Kutus site had the lowest (Table 2, Supplemental Fig. S1). The RILs were significantly (*P* < 0.05) different for PWPP (Table 2). Similarly, there were no significant differences (*P* < 0.05) for pods per plant (PPP) between the two parents in all the three sites. However, the RILs had significant (*P* < 0.001) difference for PPP across the three sites (Table 2). The mean value for the RILs across the sites varied from 0.25-149.30. The Mariira farm site had the highest number of PPP (43.51) while the Kutus site (26.77) had the lowest (Supplemental Fig. S1).

**TABLE 2.** Means of pod traits for snap bean parents and RILs evaluated in three sites in Kenya.

| Site/Trait | Parents |  | t-test | RILs (n=184) |  |  |
| --- | --- | --- | --- | --- | --- | --- |
|  | Vanilla | MCM5001 |  | Mean±SE | Range | ANOVA |
| **Kutus** |  |  |  |  |  |  |
| PWPP<sup>a</sup> (g) | 57.30±20.15 | 57.92±31.05 | ns | 49.00±13.22 | 0.54-152.50 | *** |
| PPP | 28.63±8.41 | 31.38±15.44 | ns | 26.77±6.44 | 0.40-83.00 | *** |
| PLF (cm) | 12.24±0.63 | 8.10±0.30 | * | 9.71±0.36 | 6.24-12.78 | *** |
| PD (mm) | 6.77±0.06 | 8.44±0.06 | ** | 7.41±0.14 | 6.50-8.79 | *** |
| PSFP | 0.36±0.07 | 0.77±0.23 | ** | 0.46±0.09 | 0.07-1.00 | *** |
| PFFP | 0.00±0.00 | 2.00±0.00 | *** | 1.08±0.20 | 0.00-2.00 | *** |
| PSDP | - | - | - | - | - |  |
| PFDP | - | - | - | - | - |  |
| **Don Bosco** |  |  |  |  |  |  |
| PWPP (g) | 88.79±19.79 | 103.39±21.56 | ns | 89.93±18.71 | 3.66-291.60 | *** |
| PPP | 38.04±6.45 | 43.99±8.21 | ns | 41.01±7.94 | 1.9-122.80 | *** |
| PLF (cm) | 13.51±0.57 | 9.33±0.14 | * | 10.37±0.33 | 7.56-15.24 | *** |
| PD (mm) | 7.16±0.06 | 8.80±0.07 | ** | 7.91±0.12 | 6.69-9.00 | *** |
| PSFP | 0.23±0.04 | 1.00±0.00 | ** | 0.75±0.12 | 0.11-1.00 | *** |
| PFFP | 0.00±0.00 | 2.00±0.00 | *** | 1.13±0.37 | 0.00-2.00 | *** |
| PSDP | 0.00±0.00 | 10.00±0.00 | *** | 5.87±2.09 | 0.00-10.00 | *** |
| PFDP | 0±0.00 | 10±0.00 | *** | 4.41±1.98 | 0.00-10.00 | *** |
| **Murang'a** |  |  |  |  |  |  |
| PWPP (g) | 81.48±18.05 | 95.61±46.55 | ns | 82.28±27.76 | 0.63-301.2 | * |
| PPP | 42.93±9.97 | 53.74±23.78 | ns | 43.51±13.44 | 0.25-149.3 | ** |
| PLF (cm) | 13.32±0.55 | 8.03±0.18 | ** | 10.03±0.41 | 7.15-14.15 | *** |
| PD (mm) | 6.84±0.04 | 8.29±0.03 | *** | 7.35±0.13 | 6.50-8.77 | *** |
| PSFP | 0.52±0.12 | 1.00±0.00 | * | 0.60±0.13 | 0.20-1.00 | *** |
| PFFP | 0.00±0.00 | 2.00±0.00 | *** | 1.11±0.34 | 0.00-2.00 | *** |
| PSDP | 0.00±0.00 | 10.00±0.00 | *** | 6.29±1.74 | 0.00-10.00 | *** |
| PFDP | 0.00±0.00 | 10.00±0.00 | *** | 5.91±1.70 | 0.00-10.00 | *** |
| **Combined sites** |  |  |  |  |  |  |
| PWPP (g) | 75.86±10.78 | 85.64±18.68 | ns | 73.7±22.08 | 0.54-301.2 | *** |
| PPP | 36.53±4.70 | 43.03±9.11 | ns | 37.07±10.41 | 0.25-149.30 | *** |
| PLF (cm) | 13.02±0.35 | 8.49±0.24 | *** | 10.01±0.38 | 6.24-15.24 | *** |
| PD (mm) | 6.92±0.07 | 8.51±0.08 | *** | 7.56±0.13 | 6.50-9.00 | *** |
| PSFP | 0.37±0.06 | 0.92±0.08 | *** | 0.60±0.12 | 0.07-1.00 | *** |
| PFFP | 0.00±0.00 | 2.00±0.00 | *** | 1.11±0.31 | 0.00-2.00 | *** |
| PSDP | 0.00±0.00 | 10.00±0.00 | *** | 6.06±1.92 | 0.00-10.00 | *** |
| PFDP | 0.00±0.00 | 10.00±0.00 | *** | 5.17±1.83 | 0.00-10.00 | *** |

PWPP<sup>a</sup>-pod weight per plant, PPP-pods per plant, PLF-pod length, PD-pod diameter, PSFP-pod string fresh pods, PFFP-pod fiber fresh pods, PSDP-pod string dry pods, PFDP-pod fiber dry pods, \*, \*\*, \*\*\* and ns, significant at p<0.05, 0.01, 0.001 and not significant respectively, t-test represent the level of significance for the p-value of a t-test between parental means, ANOVA represents the level of significance among RILs.

#### Pod dimensions

There were significant (*P* < 0.05) differences between the two parents and the RILs for pod length in the field (PLF) across the three sites. Vanilla had longer pods than MCM 5001 across all sites. The RILs’ mean PLF were 9.71, 10.37, and 10.03 cm at the Kutus, Don Bosco, and Mariira farms, respectively. The average PLF across the sites was 10.01 cm (Table 2). The Don Bosco site had the longest (10.37cm) PLF values while the Kutus site had the shortest (9.71 cm) (Supplemental Fig. S1). Results for pod diameter (PD) showed significant (*P* < 0.05) differences between the parents and the RILs in all three sites. The mean PD for the parents and RILs was highest at the Don Bosco site. The RIL population means for pod diameter were 7.41, 7.91 and 7.35 mm for the Kutus, Don Bosco and Mariira farms, respectively (Table 2, Supplemental Fig. S1).

#### Pod string

Significant differences (*P* < 0.05) were recorded between the two parents for pod string in both fresh (PSFP) and dry pods (PSDP). The RIL population mean for PSFP was 0.60 and for PSDP was 6.06 (Table 2). The Kutus site had the lowest PSFP (0.46), while Don Bosco had the highest (0.75). RILs were significantly (*P* < 0.001) different for PSFP and PSDP in all the sites (Table 2, Supplemental Fig. S1).

#### Pod fiber

Significant differences (*P*<0.001) were detected for both PFFP and PFDP for all the three sites. PFFP ranged from 1.08 (Kutus) to 1.13 (Don Bosco). Significant (*P* < 0.001) differences for PFFP and PFDP of RILs were observed in all the three sites. The RIL population mean PFFP was 1.11 and for PFDP was 5.17 (Table 2, Supplemental Fig. S1).

### Phenotypic correlation analysis

Highly significant correlations (*P* < 0.001) were identified between numerous measured traits (Fig. 1). This included correlations among related traits, such as string traits (PSSS, PSS, PSFP, and PSDP), which had Pearson correlation coefficients ranging from *r*=0.41 (PSSS vs. PSFP) to *r* = 0.96 (PSSS vs. PSS). Wall fiber traits (PFS, PWF, PFFP, and PFDP) had Pearson correlation coefficients ranging from *r* = 0.41 (PFDP vs. PWF) to *r* = 0.90 (PFS vs PWF). The two productivity traits, pod weight per plant (PWPP) and pods per plant (PPP), were highly correlated (*r* = 0.93).

**FIGURE 1.**
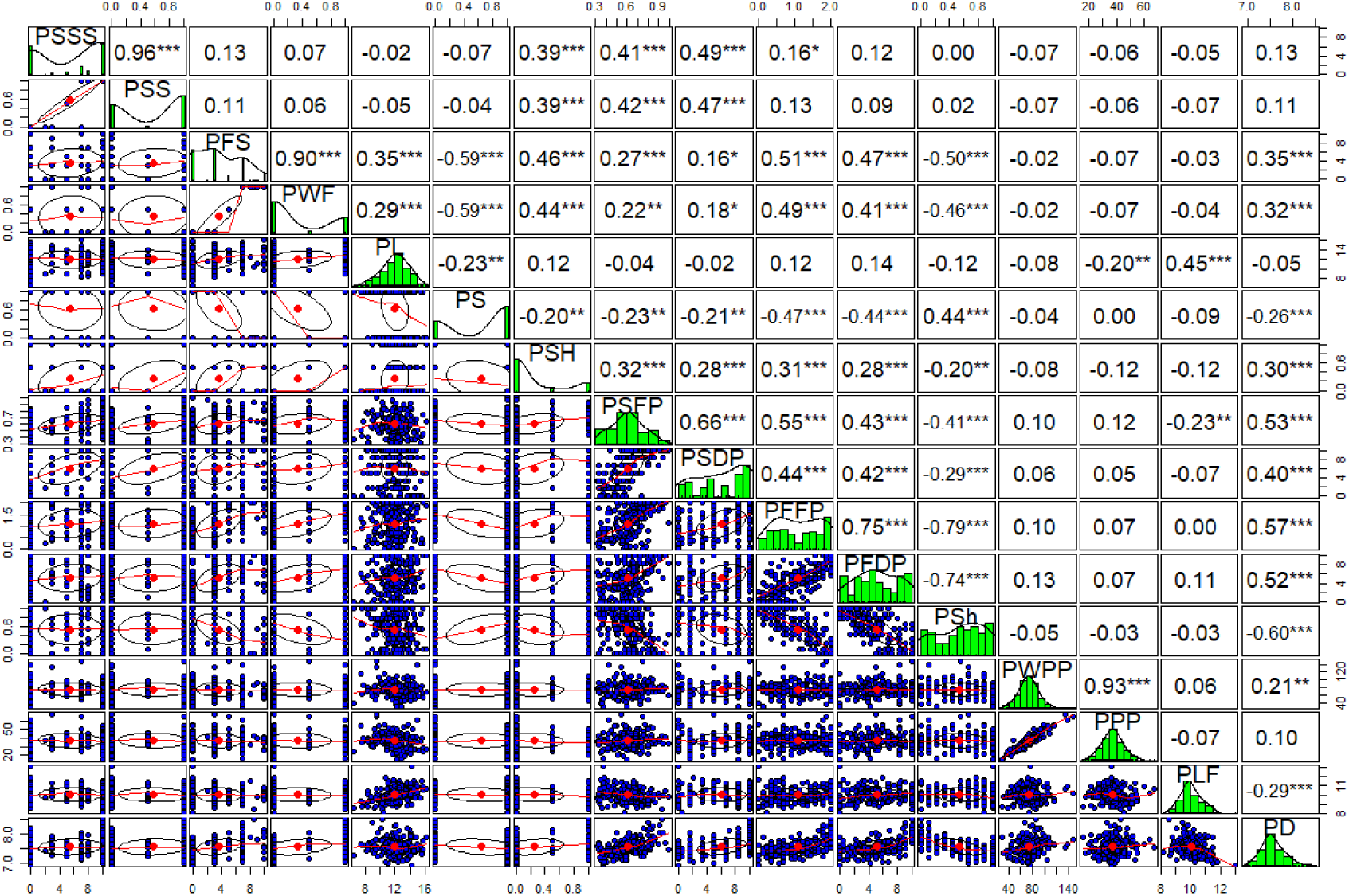
Phenotypic correlations between various measured traits pod quality and yield traits of snap beans. Upper panels indicate Pearson correlation coefficients (*r*), while diagonal and lower panels represent distributions of the data among RILs. Abbreviations: Pod suture string scale (PSSS); Pod suture string (PSS); Pod fiber scale (PFS); Pod wall fiber (PWF); Pod length (PL); Pod shape (PS); Pod shattering (PSH); Pod string fresh pods (PSFP); Pod string dry pods (PSDP); Pod fiber fresh pods (PFFP); Pod fiber dry pods (PFDP); Pod shape (PSh); Pod weight per plant (PWPP); Pods per plant (PPP); Pod length (PLF); Pod diameter (PD).

Highly significant correlations (P < 0.001) were also identified between trait categories, such as between pod cross sectional shape traits (PS and PSh) and pod wall fiber traits. Correlations between these ranged from *r* = –0.44 (PFDP vs. PS) to *r* = –0.79 (PFFP vs. PSh). The two productivity traits PPP and PWPP varied independently from all quality traits (P < 0.05), except for a single significant correlation (*P* < 0.01) between pods per plant (PPP) and pod length (PL) (*r* = –0.20) (Fig. 1).

Simultaneously, several other traits were uncorrelated in the population, including pod shape (PS) vs. pods per plant (PPP) (*r* = 0.00, *P* > 0.05); and pod suture string scale (PSSS) vs. pod shape (PS) (*r* = 0.00, *P* > 0.05) (Fig. 1).

### Vanilla x MCM 5001 genetic map

The genetic map covered 1952 cM, with an average marker density of 3 SNPs per cM. The linkage group size varied from 139 cM (Pv10) to 238 cM (Pv02) with an average size of 177.54 cM. The number of SNPs per chromosome varied from 180 (Pv05) to 786 (Pv01) with an average number of 541 SNPs per linkage group (Table 3).

**TABLE 3.** Number of SNPs and linkage length of each chromosome in the Vanilla x MCM 5001 genetic map.

| Chromosome number | SNPs | cM | Marker density<br>(markers/cM) |
| --- | --- | --- | --- |
| Pv01 | 786 | 181 | 4.4 |
| Pv02 | 469 | 238 | 2.0 |
| Pv03 | 692 | 194 | 3.6 |
| Pv04 | 326 | 162 | 2.0 |
| Pv05 | 180 | 139 | 1.3 |
| Pv06 | 370 | 144 | 2.6 |
| Pv07 | 484 | 184 | 2.6 |
| Pv08 | 710 | 246 | 2.9 |
| Pv09 | 570 | 178 | 3.2 |
| Pv10 | 645 | 138 | 4.7 |
| Pv11 | 719 | 149 | 4.8 |
| Total | **5951** | **1952** | **3.1** |

### QTL analysis

A total of 44 QTLs for PWPP, PPP, PL, PD, PS, PF, PSh, and PSH were identified for all pod traits that were evaluated (Table 4). The QTLs were distributed on seven (Pv01, Pv02, Pv03, Pv04, Pv06, Pv07 and Pv08) of the eleven chromosomes of common bean (Fig 2, Table 4). Maximum LOD scores by trait varied from 2.82 for PWPP to 38.02 for PSS. The phenotypic variation explained by the identified QTLs varied from 6.81% for PWPP to 61.39% for PSS. Vanilla was the donor of favourable QTL alleles for all pod traits except PWPP and PPP.

**FIGURE 2.**
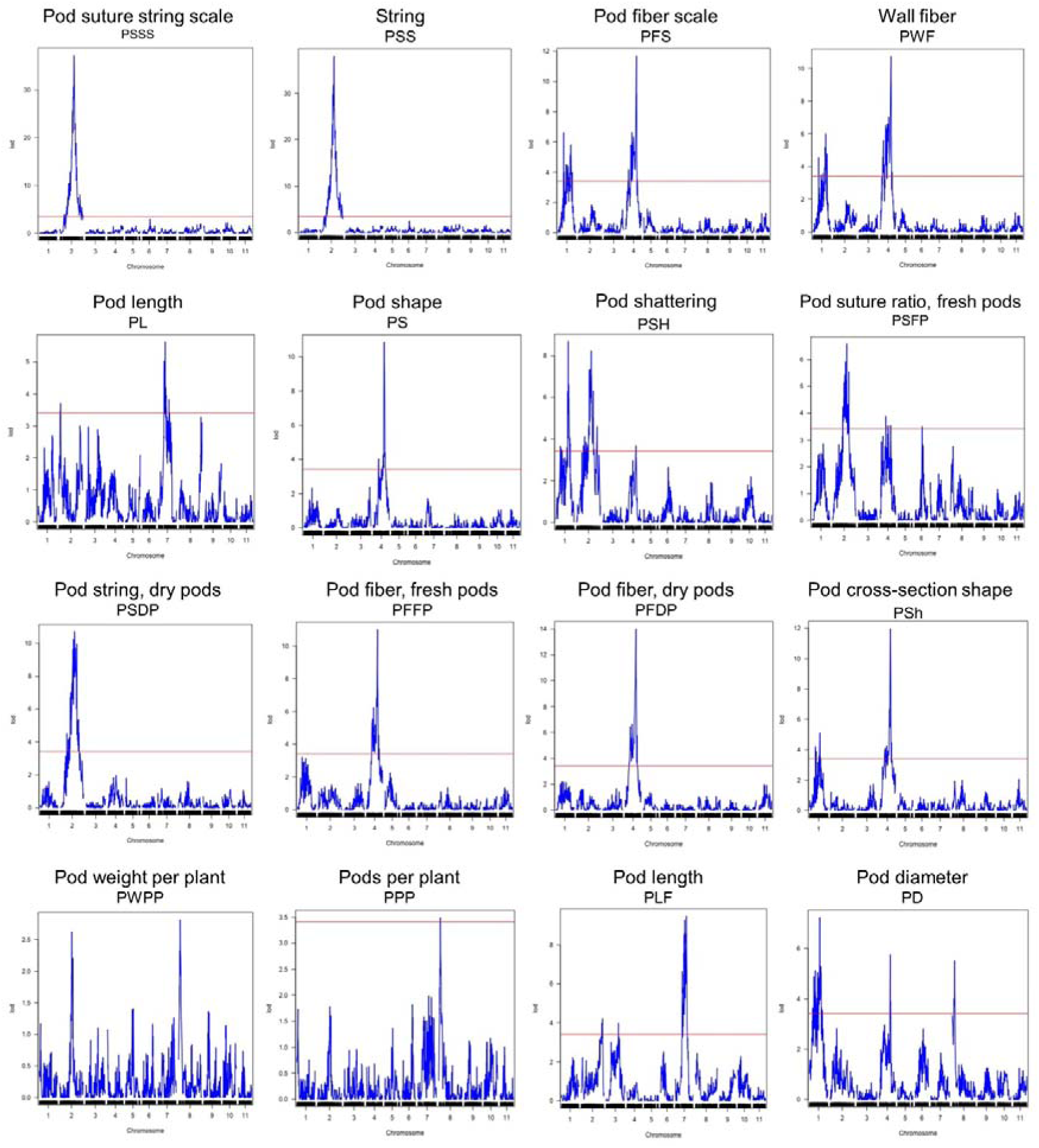
QTL plots for the sixteen pod quality and yield-related traits. The horizontal axis indicates the chromosomes and the vertical axis indicates the LOD score. The dashed line indicates the significance threshold at *P* = 0.05 based on 1000 randomized data permutations.

**TABLE 4.**
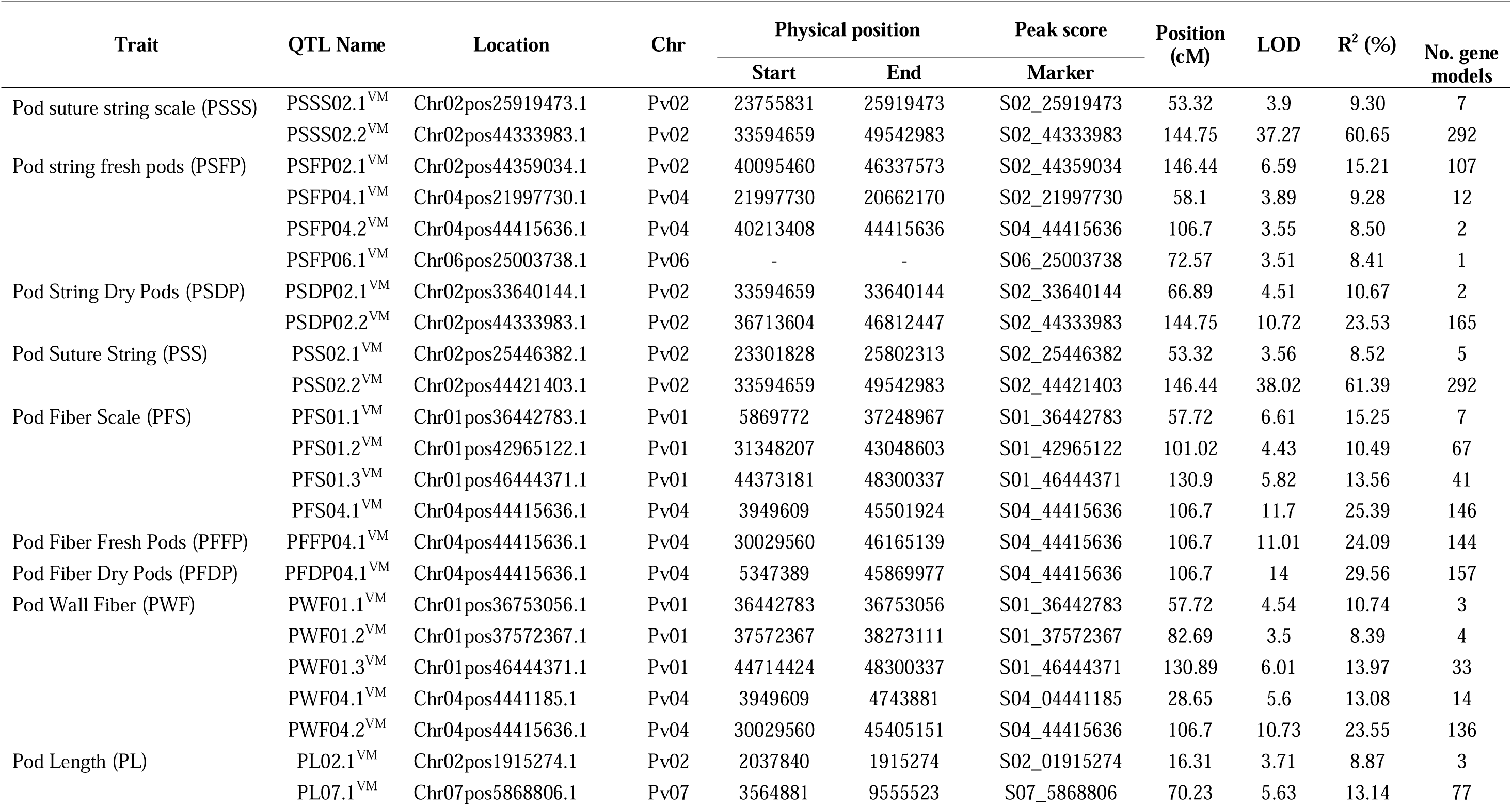
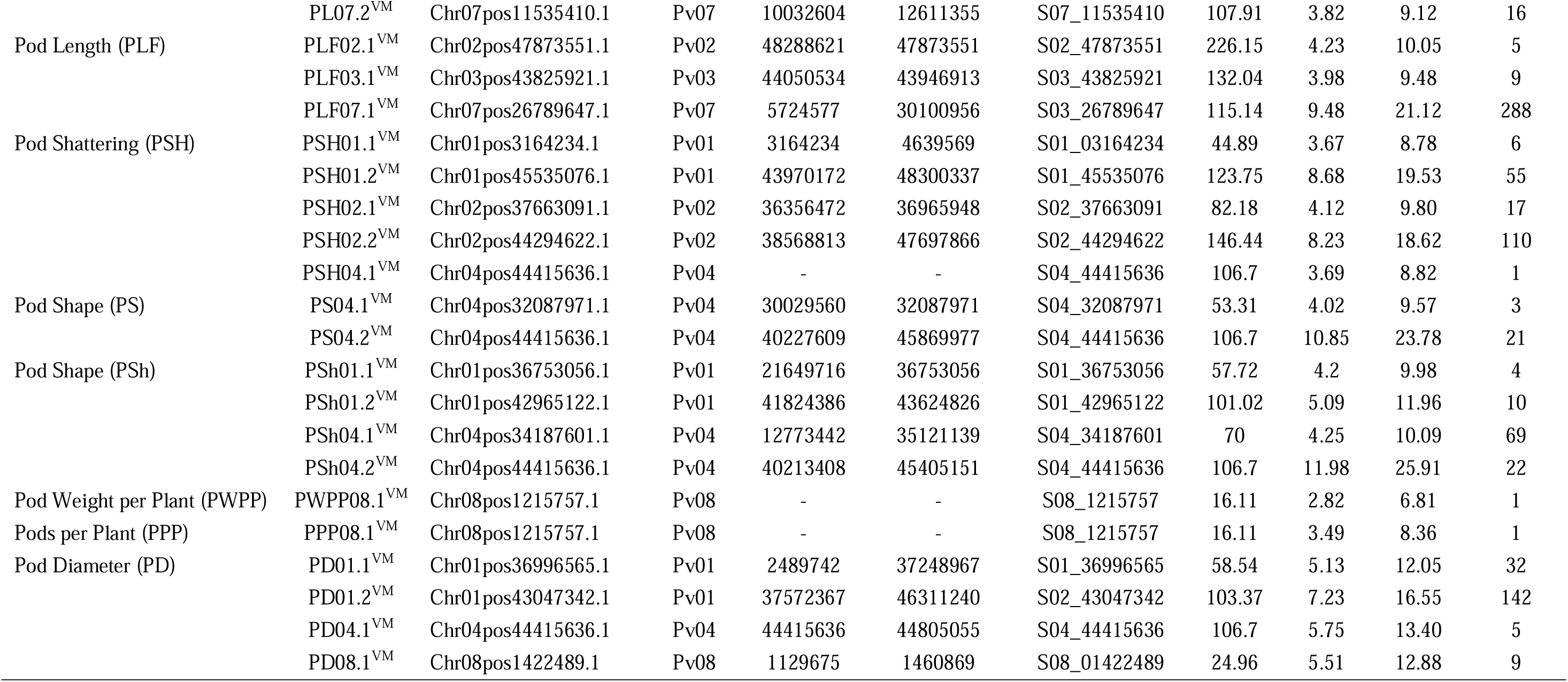
Quantitative trait loci identified using 184 recombinant inbred lines of Vanilla x MCM 5001 evaluated for pod quality and yield traits in three sites.

#### Pod weight per plant

The QTL with the highest LOD score for PWPP was detected on Pv08. The QTL explained 6.81% of the phenotypic variation but was not significant (LOD=2.82) based on the LOD threshold of 3.413. MCM5001 contributed a positive allele effect at PWPP08^VM^.

#### Number of pods per plant

QTL analysis detected one QTL on Pv08 (PPP08.1^VM^). The QTL accounted for 8.4% of the phenotypic variation. This QTL (LOD = 3.49) was just above the LOD threshold of 3.413for that trait and MCM5001 was the donor of the QTL allele conditioning higher phenotype values for PPP.

#### Pod length

A total of six PL QTL were identified on three chromosomes in both greenhouse and field trials (Fig 2; Table 4). Of the six QTL, three (PL02.1^VM^, PL07.1^VM^ and PL07.2^VM^) were identified in the greenhouse environment with a LOD score of 3.71-5.63. PL07.1^VM^ (3.56-9.56 Mb, *R*^2^ = 13.14%) was the most significant QTL for pod length in the greenhouse. Three PL QTL (PL02.1^VM^, PL03.1^VM^ and PL07.1^VM^) were identified in the field trials. PL02.1^VM^ and PL07.1^VM^ were major QTLs explaining phenotypic variation of 10.05% and 21.12% respectively. The phenotypic variation explained varied from 8.87% to 21.12% for both greenhouse and field trials and MCM5001 introduced alleles that lead to a negative effect on PL.

#### Pod diameter

A total of four PD QTLs were identified on chromosome Pv01, Pv04, and Pv08. PD01.2^VM^ and PD04.1^VM^ were the most significant PD QTLs explaining phenotypic variation of 16.55% and 13.40%, respectively (Table 4). PD04.1^VM^ (44.415 Mb) overlapped with a QTL for pod shape in the greenhouse (PS04.2^VM^), fresh pod string in the field (PSFP04.2^VM^), fresh pod fiber (PFS04.1^VM^, PFFP04.1^VM^, and PWF04.2^VM^), pod fiber dry pods in the field (PFDP04.1^VM^), and pod shattering (PSH04.1^VM^) (Table 4). MCM5001 donor produced flat pods with a broad cross-section and hence conditioning higher phenotype values for PD.

#### Pod string

A combined total of ten pod string QTL were identified in greenhouse and field trials on chromosome Pv02, Pv04 and Pv06. Eighty percent of the QTLs were located on chromosome Pv02. PSSS02.2^VM^ and PSS02.2^VM^ were the most significant QTLs explaining phenotypic variation of 60.65% and 61.39%, respectively (Table 4). Both PSSS02.2^VM^ and PSS02.3^VM^ QTL spanned from 33,594,659 to 49,542,983 bp. Five of the PS QTLs had major effect explaining more than 10% phenotypic variation. PSFP04.2^VM^ explained 8.50% of the variation for fresh pod string, overlapped with major loci for PFS04.1^VM^, PFFP04.1^VM^, PFDP04.1^VM^, PWF04.2^VM^, PSH04.1^VM^, PS04.2^VM^, PD04.1^VM^ and PSh04.2^VM^. MCM5001 alleles contributed to increased PS.

#### Pod fiber

Eleven QTL related to PF were detected on two chromosomes (Pv01 and Pv04) in greenhouse and field trials for fresh and dry pods. Major effect QTLs controlling PF were detected in a 106.7 cM region on Pv04 (Fig 2; Table 4). PFS04.1^VM^, PFFP04.1^VM^, PFDP04.1^VM^ and PWF04.2^VM^ were the most significant PF QTL explaining 25.39%, 24.09%, 29.56% and 23.55%, of the phenotypic variation respectively. In chromosome Pv01, PWF01.3^VM^ and PFS01.1^VM^, explained 13.97% and 13.56% phenotypic variation, respectively.

Six QTLs for pod shape were detected in two chromosomes (Pv01 and Pv04) in both greenhouse and field trials. The QTL with the highest contribution to the trait (PS04.2^VM^, greenhouse and PSh04.2^VM^, field) were located on Pv04 and explained 23.78% and 25.91% of the phenotypic variation, respectively (Table 4). MCM5001 provided alleles that increase phenotype values for pod fiber, while reducing the pod shape (increased flatness).

#### Pod shattering

Pod shattering is an important trait associated with seed dispersal, which was modified during domestication from the wild dehiscent (shattering) state to pod indehiscence (non-shattering). QTL analysis detected five PSH QTL on three chromosomes (Pv01, Pv02, and Pv04). PSH01.2^VM^ and PSH02.2^VM^ were the most significant QTL for the trait explaining a phenotypic variation of 19.53% and 18.62%, respectively (Table 4). PSH04.1^VM^ on Pv04 explained 8.82% of the phenotypic variation and co-located with QTLs for pod wall fiber (PFDP04.1^VM^, PFS04.1^VM^, PFFP04.1^VM^, and PWF04.2^VM^), pod shape (PS04.2^VM^ and PSh04.2^VM^) and pod string (PSFP04.2^VM^). MCM5001 contributed QTL alleles that condition higher phenotypic values for PSH.

## DISCUSSION

The utilization of genetic diversity is fundamental for the efficient identification of superior genotypes across all crops, including snap beans. Enhancing crop quality relies on the extent of genetic variation observed for economically significant traits. Consequently, the assessment and exploitation of genetic diversity toward desired objectives play a pivotal role in any endeavour aimed at enhancing crop yields (Singh et al., 2018). The use of a mapping population with parental lines that show extreme and contrasting phenotypes is an important resource for unravelling the main genetic architecture involved in snap bean pod characteristics. Snap bean cultivars exhibit slender, elongated, cylindrical pods with markedly diminished fiber content, alongside thickened pod walls and diminutive seeds (Singh and Singh, 2015).

Breeding for pod quality and yield traits is a major objective for snap bean improvement programs (Singh and Singh, 2015; García-Fernández et al., 2022). Pod quality traits influence consumers’ preferences and palatability, while pod yield related traits influence farmers adoption of a new variety (Yuste-Lisbona et al., 2014; Singh et al., 2018; García-Fernández et al., 2024). The present study utilised a RIL population originating from the hybridization of two parents with contrasting pod phenotypes to locate the position of pod quality and yield QTLs. The parents and the RILs were significantly different (p < 0.05) for all the traits and sites apart from PWPP and PPP. This indicated that genetic variation existed between the parents and among the RILs for the evaluated traits.

In the current study, PWPP was positively significantly correlated with other traits apart from PFDP. This indicates the potential usefulness of these traits in addition to PWPP when selecting for pod yield. PD is highly correlated to pod string and pod fiber, which are very important traits for pod quality. These results were consistent with the findings of previous studies (Hagerty et al., 2016; García-Fernández et al., 2024; García-Fernández et al., 2021).

QTL analysis was conducted to gain insight into the genetic architecture of pod quality and yield traits under different environmental conditions. A large number of identified QTLs for pod quality and yield traits involves a large set of genes for different pod morphological traits. High phenotypic variation (*R^2^*) explained by any single QTL suggest the effect of additive genes in the control of pod quality traits (Jusoh, 2017).

Various mapping studies in common bean have reported QTLs for pod yield and yield components on several chromosomes (Koinange et al., 1996; Tar’an et al., 2002; Beattie et al., 2003; Blair et al., 2006; Kamfwa et al., 2015). In this study, pod weight and number of pods per plant were positively correlated and located on chromosome Pv08, in agreement with Kamfwa et al. (2015) who identified significant SNPs for pod weight per plant on Pv08 in the Andean Diversity Panel (ADP) of common bean (Cichy et al., 2015). Further, Koinange et al. (1996) reported QTL for number of pods per plant on Pv01 and Pv08 from an inter-gene pool cross of Midas (cultivated wax snap bean) × G12873 (wild Mesoamerican accession). Additional QTL for pod weight have been reported on Pv02, Pv03, Pv05, Pv07, Pv09 and Pv11 (Tar’an et al., 2002, Beattie et al., 2003; Blair et al., 2006; Kamfwa et al., 2015) when studied in other populations.

The reduction of pod suture string is crucial for pod edibility and, therefore, constitutes a distinguishing characteristic between dry and snap beans (Parker et al., 2022). Ten QTLs were identified for pod string on Pv02, Pv04, and Pv06. Our most significant QTL for string trait in both trials were found on Pv02 in the direct vicinity of *PvIND* (*Phvul.002G271000*), which was recently shown to be the major factor in pod string formation due to gene duplication, retrotransposon insertion, and gene overexpression (Parker et al., 2022). Our Pv02 mapping results agree with those of Koinange et al. (1996), who first reported one major pod string locus (*St*) on Pv02. Furthermore, Gioia et al. (2013) first mapped the gene *PvIND* near the *St* locus. Linkage analysis by Hagerty et al. (2016) and Arkwazee et al. (2022) fine-mapped the *St* locus in a 0.5 Mb region on Pv02. The present study identified a string QTL that is unique to fresh pods on Pv06 (PSFP06.1^VM^). These results may match a QTL identified by Davis et al. (2006), who mapped strings to Pv06, and García-Fernández et al. (2024), who reported a Pv06 pod edibility QTL. PSFP06.1^VM^ in this study mapped to 25 Mb, while *EDIBILITY6^TUM^* spanned positions 15.9 to 25.7 Mb on chromosome Pv06 (García-Fernández et al., 2024). PSFP04.2^VM^ overlapped with major loci for pod wall fiber, pod diameter and pod shattering indicating the key role of the positions in the genetic control of these traits. This is the first report of a pod string QTL on Pv04.

Pod wall fiber is one of the most important agronomic traits, which plays a role in edibility and influencing consumers’ preference for snap bean. Eleven QTLs for pod wall fiber were identified on Pv01 and Pv04. Koinange et al. (1996) mapped pod wall fiber on Pv02 being conditioned by a single gene and co-segregating with pod suture string. On chromosome Pv01, pod wall fiber co-localized with pod shattering, pod diameter, and pod shape while on chromosome Pv04 it co-localized with pod string, pod shattering, pod diameter and pod shape. Hagerty et al. (2016) mapped pod fiber on Pv04 and pod string on Pv02, indicating that they have independent genetic control. Pod wall fiber, pod shape, and pod diameter were phenotypically correlated in this study. Our results are in agreement with Hagerty et al. (2016), who found significant correlations between pod wall fiber, pod width, and pod height. QTLs for pod wall fiber, pod shape, and pod diameter clustered in both Pv01 and Pv04.

Our mapping of pod shattering identified five QTLs none of which have been previously mapped, but which correspond to the unique pod traits of snap beans, such as loss of wall fiber (Pv01 and Pv04) and pod string (Pv02). Low levels of fiber deposition in the pod wall and pod suture strings in snap beans are correlated with extreme resistance to pod shattering (Parker et al., 2021a). Previous research has identified QTL for bean pod shattering on Pv02, Pv03, Pv04, Pv05, Pv08, and Pv09 (Koinange et al., 1996; Rau et al., 2019; Parker et al., 2020). Candidate genes within each of these genomic regions have been proposed, including *PvIND* (Pv02), the major suture string factor; *PvPdh1* (Pv03), a major locus controlling pod wall fiber torsion and pod shattering in common bean; MYB family transcription factors, such as *PvMYB26* and *PvMYB46* (Pv05 and Pv08), WRKY family transcription factors (Pv08), polygalacturonases (Pv08 and Pv09), and cellulose synthase (CESA7 on Pv09) (Gioia et al., 2013; Rau et al., 2019; Parker et al., 2020; Parker et al., 2021b; Di Vittori et al., 2021). This study mapped five QTL for pod shattering in three chromosomes (Pv01, Pv02 and Pv04). The most significant QTL on Pv01 (PSH01.2^VM^) spanned the 43,970,172-48,300,337 bp region, which is a novel QTL in this study. The shattering loci identified in this study on Pv01, Pv02, and Pv04 are co-located with QTLs we identified for pod wall fiber and pod string. For example, PSH01.2^VM^ corresponded to the wall fiber QTLs PWF01.3^VM^ and PFS01.3^VM^; PSH02.2^VM^ corresponded to the most significant QTLs for the four pod string traits (PSS02.2^VM^, PSSS02.2^VM^, PSDP02.2^VM^, and PSFP02.1^VM^). Additionally, the most significant SNP for shattering on Pv04 (PSH04.1^VM^) corresponded to the most significant SNP for each of the four wall fiber traits (PFDP04.1^VM^, PFS04.1^VM^, PFFP04.1^VM^, and PWF04.2^VM^), as well as the most significant SNP for pod shape (PS04.2^VM^). Our results reinforce the notion that pod string and wall fiber are crucial for pod shattering in common bean, which had not previously been demonstrated empirically.

Pod length was mapped on Pv02, Pv03, and Pv07, for a combined six QTLs in the greenhouse and field. Koinange et al. (1996) analyzed pod length as a measure of the increase in size of the harvested organs in domesticated common beans compared with their wild progenitors, and mapped this to Pv01, Pv02, and Pv07, explaining 37% of the phenotypic variation. The most significant QTL for pod length in this study was on Pv07 (PLF07.1^VM^) explaining 21% of the phenotypic variation for the trait. Pod length showed a positive phenotypic correlation with pod weight per plant and number of pods per plant, although pod length was not linked to those traits because the QTL for pod length were found on Pv02 and Pv07 whereas pod weight per plant and pods per plant were observed on Pv08. Further, there were no significant correlation between pod length with pod diameter, pod string, and pod wall fiber. Therefore, in case larger or smaller sieve size bean is desired by a breeding program, this could be achieved independently of pod length.

As one of the major pod quality traits, pod diameter is important in snap bean breeding. Breeders require an understanding of the relationship between desirable processing traits to breed commercially acceptable snap bean varieties. The QTL clustering of pod wall fiber, pod string, pod shape, pod shattering, and pod diameter traits in close proximity on Pv01, Pv02, and Pv04 make these particularly useful chromosomal regions for breeders selecting for new fresh and canning market snap beans. Given the proximity and correlation of these traits, we suggest that QTL for pod wall fiber, pod string, pod shape, pod shattering, and pod diameter may be pleiotropically controlled by one or a small number of genes on each of these chromosomes. The most significant QTL for pod wall fiber and pod shape traits consistently identified position 44,415,636 on Pv04 as the most significant locus.

Our QTL mapping for pod shattering were co-located with major QTLs for pod wall fiber and pod string. Pod shattering, a significant characteristic linked to seed dispersal, underwent a change to indehiscent pods during the process of domestication. Our results contrast with other recent research, which identified shattering QTL located on chromosomes Pv03, Pv05, Pv08, and Pv09 (Rau et al., 2019; Parker et al., 2020). Legume pod dehiscence is known to be controlled by downstream NAC and MYB family transcription factors which also regulate the formation of secondary cell walls (Nakano et al., 2015; Ohtani and Demura, 2019; Takahashi et al., 2020; Watcharatpong et al., 2020; Zhang and Singh 2020; Chen et al., 2021; Gupta et al., 2021). Additional genes responsible for controlling legume shattering, such as the soybean dirigent gene *PDH1* and *PvPdh1* in common bean, influence the torsion of pod valves without causing anatomical alterations, as observed in studies by Suzuki et al. (2009) and Parker et al. (2020). However, these genes are not expected to regulate the formation of pod suture strings. These QTL were not significantly associated with pod shattering in our population. The parents of our population were of Andean (Vanilla) and Middle American (MCM5001) origin, and therefore came from independent domestication events. The lack of shattering QTLs at the candidate genes on Pv03 (*PvPdh1*) and Pv05 (*MYB26*) indicate that the parents each likely carry alleles with similar phenotypic effects at each gene (e.g., both parents with loss-of-function alleles, or both with fully-functional alleles). This is an indication that orthologous pathways may have been selected in each independently domesticated gene pool. Our work, unlike previous shattering studies, was based on a population developed without backcrossing with a snap bean parent. Our results strongly suggest that the selection for snap bean pod quality traits, such as loss of pod wall fiber and pod strings, has pleiotropically greatly reduced the levels of pod shattering in these materials. These genes may therefore be of value for breeding dry bean for shattering resistance in arid conditions (Parker et al. 2020; 2021).

Several candidate gene models were identified and/or supported through our analyses. Many of these were also collocated, likely as the result of pleiotropy The most significant QTL for pod wall fiber and pod shape traits consistently identified position 44,415,636 on Pv04 as the most significant locus. This SNP is found within the gene model *Phvul.004G143500*, which encodes a homeobox-leucine zipper protein. In Arabidopsis, the closest orthologs of this gene include *HOMEODOMAIN GLABROUS 11* and *12* (*HDG11* and *HDG12*; *AT1G73360* and *AT1G17920* respectively), and *PROTODERMAL FACTOR 2* (*PDF2*, *AT4G04890*). In Arabidopsis, these genes are involved in the maintenance of floral organ identity, cell wall identity, and patterning of outer layers of plant organs (Nakamura et al., 2006; Kamata et al., 2013).

While all pod string evaluations identified the *St* gene on Pv02, secondary QTL for pod string were only identified in the warm Kenyan field conditions, but not in the cool winter greenhouse conditions of Davis, CA, USA. These QTL, on Pv04 and Pv06, may therefore underlie the *Temperature-sensitive* (*Ts*) partial string locus of Drijfhout (1970; 1978). The chromosome Pv06 QTL for pod suture ratio in fresh pods was identified on Pv06 at 25,003,738. The common bean homolog of *SHOOTMERISTEMLESS, PvSTM (Phvul.006G145800)*, is found at 25.18 Mb on the same chromosome. In Arabidopsis, *STM* is an important regulator of floral patterning and development of the replum (Girin et al., 2009), which is partly homologous to the suture fiber bundle of common bean. *PvSTM* is, therefore, a potential candidate for the control of partial pod string. Another Pv06 candidate is *Phvul.006G144300*, at 25.02 Mb, which is homologous with SHOU4-like genes in Arabidopsis that regulate cellulose synthesis by controlling exocytosis of cellulose synthase enzymes. The most significant Pv06 QTLs for traits such as pod diameter, pod shattering, string, and string scale were also found in this region, specifically at 25,003,738 bp and 24,899,534 bp. This indicates that the locus may pleiotropically affect other pod traits.

A region at the beginning of Pv08 was significantly associated with pod number per plant and pod diameter. *Phvul.008G014000* is found near the QTL peak for pod number per plant and encodes a WUSCHEL-related homeobox (WOX) 10-related protein. WOX genes are known to regulate meristematic stem cells and floral patterning (van der Graaff et al., 2009). Whether pod diameter and pod number are in linkage or are pleiotropically related is unclear.

Pv01 included a region significantly related to pod wall fiber, pod diameter, and pod shattering. Many of these traits are also controlled by a region on chromosome Pv04, where a candidate gene for the traits is a patterning gene responsible for specifying external surfaces such as the epidermis. Intriguingly, our Pv01 locus maps near *Phvul.001G173700* (43.0 Mb on Pv01). *Phvul.001G173700* encodes a close relative of *TRANSPARENT TESTA GLABRA 2 (WRKY44)*, a patterning gene that is also implicated in the specification of epidermis and other superficial tissues. Since wall fiber only forms on the interior surface of pods, it is possible that over-expression of these epidermal genes might reduce the expression of *MYB26* and other genes responsible for developing interior fiber cells. This pattern has been documented in Arabidopsis (Li et al., 2007).

## CONCLUSION

This study identified QTL for important pod quality and yield traits. The majority of the QTLs that were identified in this study are consistent with previous studies that used bi-parental linkage mapping with different marker resolution. Additionally, we identified novel QTLs for several pod quality and yield traits, which resulted to identification of candidate genes for pod morphological characteristics in snap bean. The identified QTLs could potentially be used as candidates for marker-assisted selection, to enhance gains in breeding for pod quality and yield traits in snap beans. Further studies using other different populations at the significant SNP loci may be necessary to validate the QTLs and their usefulness in snap bean breeding.

## ACKNOWLEDGEMENTS AND FUNDING

The authors are grateful to the Kirkhouse Trust for financial support and to the University of California, Davis, and University of Embu for provision of research facilities. Travis Parker was also partly supported by USDA-NIFA-AFRI award number 2023-67013-40001.

## CONFLICTS OF INTEREST

The authors declare that the research was conducted in the absence of any commercial or financial relationships that could be construed as a potential conflict of interest.

## AUTHOR CONTRIBUTIONS

Conceptualization: EEA. Population development: SNN. Phenotyping: SNN. Genotyping: TAP, SNN, JD. QTL mapping and analysis: TAP, SNN. Advising and administration: TAP, JD, PG, EEA. Initial draft preparation: SNN, EEA, TAP. All authors have read and agree to the final version of the manuscript.

## Notes

### Competing Interest Statement

The authors have declared no competing interest.

