## Supplemental Figure 1 for "QTL Mapping for Pod Quality and Yield Traits in Snap Bean (*Phaseolus vulgaris* L.)"

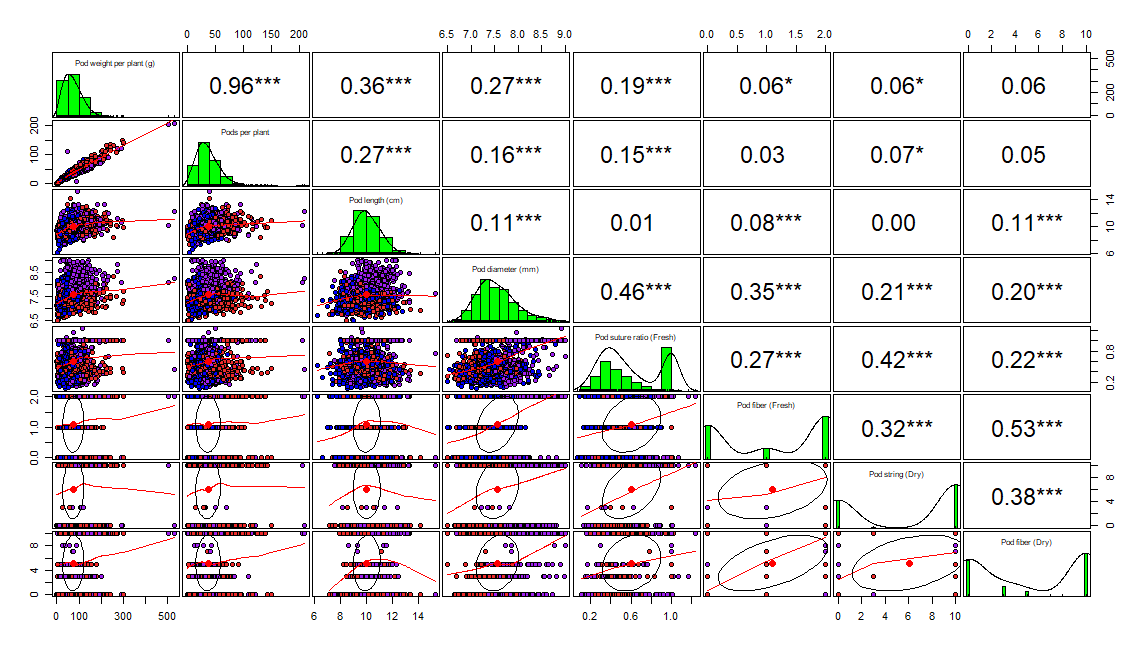

SUPPLEMENTAL FIGURE S1. Phenotypic correlations between field data for all RILs at all field sites. Upper panels indicate Pearson correlation coefficients (*r*), while diagonal and lower panels represent distributions of the data among RILs. Point color represents location: Don Bosco (purple), Mariira (red), Kutus (blue).
